# Local synaptic competition and global homeostatic regulation of synaptic resources as a unified framework for synaptic scaling

**DOI:** 10.1101/2025.04.10.648137

**Authors:** Petros E. Vlachos, Jochen Triesch

## Abstract

Neuronal firing rates must remain stable despite continuous synaptic modifications driven by Hebbian plasticity. Two compensatory mechanisms have been implicated in this stability. Heterosynaptic plasticity, which rapidly redistributes synaptic strength among competing inputs, and synaptic scaling, which adjusts synaptic efficacies globally on slow timescales. How these interact to maintain stable activity remains unclear. Here, we propose a unified framework in which local synaptic competition for a shared resource pool, combined with slow cell-wide homeostatic regulation of the total pool size, prevents runaway synaptic dynamics and ensures firing rate homeostasis. We show analytically that this architecture guarantees stability regardless of the timescale difference between Hebbian and homeostatic plasticity, and that the ratio of sensing and regulatory timescales determines whether convergence is monotonic or oscillatory. Fitting the model to activity blockade data reveals an inverse coupling between these timescales. When competition is spatially restricted to dendritic compartments, the framework reproduces experimentally observed synaptic weight distributions and local homeostatic upscaling following denervation, suggesting that heterosynaptic competition and synaptic scaling share a common mechanistic origin.

## Introduction

Neuronal firing rates remain remarkably stable over long timescales despite variations induced by learning and experience [40, 102]. Deviations from this stability are associated with a range of neurological conditions, including epilepsy [1, 44, 63], schizophrenia [55, 56], and autism spectrum disorder [72]. The primary mechanism underlying learning and information storage is Hebbian plasticity, long-term activity-dependent modifications of synaptic strength [39, 73]. However, Hebbian plasticity is inherently unstable. It introduces a positive feedback loop that can compromise the stability of cortical circuits [63, 67, 81, 102, 111, 123]. Compensatory regulation is therefore necessary, allowing the brain to encode new information while preserving the integrity of its circuits.

Synaptic scaling is a homeostatic regulatory process observed across multiple neuron types [62, 77, 87, 103, 104]. By multiplicatively adjusting synaptic strengths [107, 115], it controls neuronal activity at a firing rate set-point while preserving the relative differences between synapses and thus stored information [102]. Synaptic scaling is triggered by persistent deviations from a homeostatic activity set-point [41, 47, 103, 113].

Detecting such persistent deviations requires the neuron to integrate its own activity over time, implying a characteristic sensing timescale [47, 97, 101, 102]. The resulting signal triggers a response via transcription, translation, and receptor trafficking [46, 47, 84, 85, 90, 92]. Specifically, it affects insertion and removal of synaptic proteins, such as neurotransmitter receptors and scaffolding molecules, whose numbers correspond to synaptic strength [53, 66], therefore adjusting synaptic efficacies. This downstream regulatory process operates on slow timescales of hours to days [40, 47, 53, 78, 103]. However, how the timescales of the activity sensing and molecular regulation jointly determine the dynamics of the homeostatic response remains poorly understood.

Another compensatory mechanism that constrains runaway Hebbian synaptic dynamics is heterosynaptic plasticity. In contrast to the input-specific changes of homosynaptic plasticity, heterosynaptic plasticity describes synaptic changes on an unstimulated pathway [21]. A fundamental manifestation of heterosynaptic plasticity is the induction of modifications in the opposite direction of homosynaptic plasticity, supporting the preservation of the total synaptic strength [15, 19, 65, 110, 112]. Such modifications naturally arise due to competition among synapses over a limited pool of synaptic resources [17, 67, 80, 100, 110]. This conservation reflects the limited availability of synaptic resources whose total number remains approximately stable under baseline conditions, particularly within local dendritic domains [7, 15, 19, 75, 83, 98]. A stationary resource pool and the resulting competition offer computational benefits such as allowing the neuron to sharpen its selectivity for the dominant patterns of the input [26, 68, 76] and supporting single-trial learning [119]. However, what determines the absolute value of the conserved total synaptic strength remains an open question. Furthermore, although heterosynaptic plasticity can prevent runaway synaptic dynamics, it is unclear whether it can ensure firing-rate homeostasis, as it remains sensitive to changes in presynaptic activity levels.

The instability problem of Hebbian plasticity is partly due to its rapid temporal evolution, which unfolds within seconds to minutes [11, 19, 30, 88, 124]. Compensatory mechanisms must operate on similar timescales to counteract destabilising dynamics [1, 21, 124]. Although it remains uncertain whether heterosynaptic plasticity can implement firing rate homeostasis, it operates on the same timescales as Hebbian plasticity, making it a well-suited candidate for preventing runaway synaptic growth [15, 21, 65]. By contrast, the slow biological timescale of synaptic scaling poses a fundamental theoretical problem. Conventional, synapse-specific models of synaptic scaling must rely on comparable timescales to those of Hebbian plasticity to ensure stability [106, 122, 123]. Even then, delays in the activity sensor lead to overcompensation and oscillatory behaviour, or enforce a strict firing rate that suppresses information-carrying activity fluctuations [121]. Moreover, despite being considered a cell-wide mechanism, multiplicative scaling can be induced locally under certain conditions [7, 43, 95, 108]. Theoretical models, which typically act on the level of single synapses, cannot account for the experimentally observed local scaling. Whether synaptic scaling acts at the level of individual synapses or at the whole neuron level remains an open question [114]. Nonetheless, theoretical work highlights the need for a cell rather than synapse-specific mechanism to avoid instabilities [99]. When homeostatic forces act at the level of the neuron, stability can be ensured independently of the timescale relationship between Hebbian and homeostatic plasticity [99]. Such stability arises naturally from a multiplicative interaction between synaptic weights and a neuron-wide homeostatic factor.

Here, we propose a simple, unified framework for synaptic scaling in which local competition among synapses for a shared pool of synaptic resources, and slow cell-wide homeostatic regulation of the total size of that pool, prevent runaway synaptic weights and ensure firing rate homeostasis. Synaptic weights in our framework are normalised such that potentiation at one synapse is accompanied by depression at others. The normalisation target is not fixed but homeostatically controlled, providing a principled answer to the open question of what determines the conserved total synaptic strength. This regulation is a cell-wide mechanism, in line with theoretical suggestions [99], and is triggered by deviations from a homeostatic target activity. We show analytically that this architecture ensures stable convergence to a firing rate set-point, and that the relationship between the timescales of activity sensor and homeostatic regulator determines the nature of this convergence. We further show that input fluctuations arising from ongoing synaptic plasticity do not destabilise the neuron at steady state. Fitting the model to experimental data on activity blockade reveals a coupling between the timescales of the activity sensor and homeostatic regulator. Furthermore, when synaptic resource competition is spatially restricted to local dendritic compartments, synaptic weight distributions more closely resemble those observed experimentally. In addition, the framework naturally reproduces the layer-specific synaptic scaling observed following entorhinal denervation [108], suggesting that locally restricted scaling is a consequence of resource redistribution rather than a locally generated synaptic scaling response. Overall, despite its simplicity, the proposed framework accounts for key unresolved features of synaptic scaling within a single, biologically grounded architecture, suggesting that the phenomena of synaptic scaling and the suppression of Hebbian instability may share a common mechanistic origin.

## Results

### A model linking heterosynaptic normalization to homeostatic scaling

We begin by considering a single neuron with *N* presynaptic inputs. Each input, *i*, is characterised by the presynaptic firing rate, *x*_*i*_, and the corresponding synaptic weight, *w*_*i*_. These weights are proportional to the number of synaptic resources, such as receptor proteins and scaffolding molecules [53, 66, 74, 77]. We assume rapid competition among synapses over these resources as a form of heterosynaptic plasticity [37, 67, 100, 112]. For a synapse to potentiate, resources are drawn from nearby synapses, leading to their heterosynaptic depression (Fig. 1a, top). We implement this as normalisation over synaptic weights, which preserves their sum constant, in line with experimental observations [15, 19, 110]. We further assume slow homeostatic regulation of the total resource pool, *ρ*, as the mechanism of synaptic scaling [100, 102]. This requires the neuron to be able to sense its own activity, for example, via intracellular Ca^2+^ levels [47, 97, 101]. Then, deviations from the target firing rate set-point drive adjustments on the total sum of synaptic resources (Fig. 1a, bottom). Such adjustments could be mediated by transcription/translation [46, 47, 85, 90, 92] or other post-translational processes [84].

**Figure 1.**
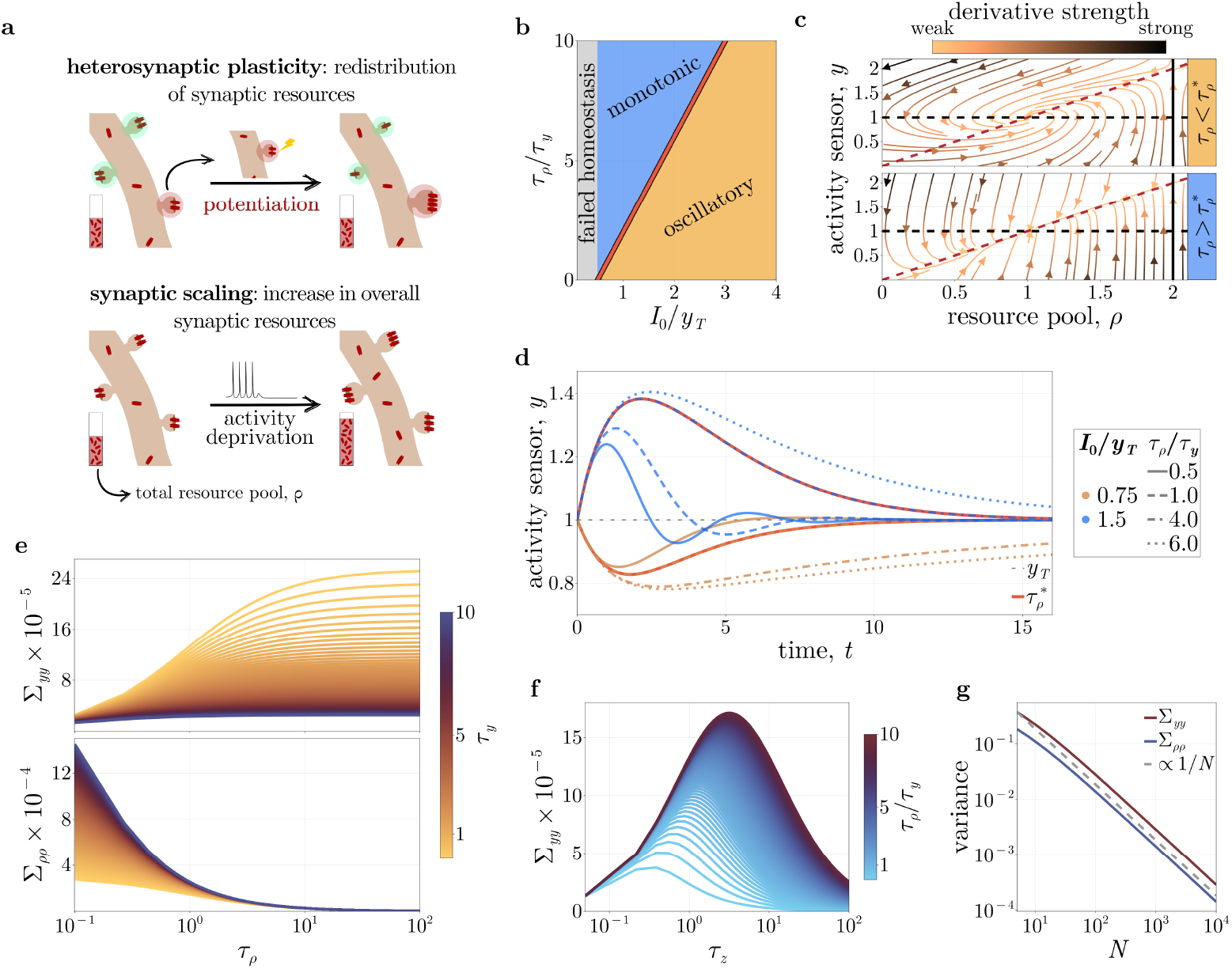
**a.** Illustration of the model. Due to competition over available synaptic resources, individual synaptic modifications induce heterosynaptic plasticity. Resources are redistributed among synapses while the total resource pool remains unchanged. At slower timescales, persistent activity changes modulate the overall resource pool. In this example, activity deprivation leads to an upregulation. **(b, c, d) Two-dimensional homeostatic model. b**. Dynamical regimes. The behaviour of the model towards the homeostatic equilibrium as a function of the ratio of the time constant of the effector, *τ*_*ρ*_, to that of the activity sensor, *τ*_*y*_, and of the input drive, *I*_0_, to the target firing rate, *y*_*T*_. The two regimes are linearly separated at the boundary condition 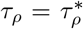 (Eq. 5). The gray area corresponds to cases where homeostasis is not possible (Eq. 4). **c**. Dynamics of the model for different timescales of the regulator, *τ*_*ρ*_, relative to the critical value 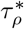. The system always converges towards the homeostatic fixed point, at the intersection of the two nullclines (dashed lines). For 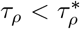, the system exhibits damped oscillations which are not present for 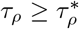. **d**. Time evolution of the activity sensor, *y*, for different magnitudes of fixed input drive, *I*_0_, and different ratios of the time constants *τ*_*ρ*_ and *τ*_*y*_. Here, the homeostatic target rate, *y*_*T*_, is set to 1 Hz. Red lines indicate the boundary between the dynamic regimes, 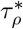. **(e, f, g) Full stochastic homeostatic model. e**. Variance of the activity sensor (top) and regulator (bottom) at the homeostatic steady state as a function of *τ*_*ρ*_ and *τ*_*y*_. Slower regulation maximises the variance of the activity sensor while that of the regulator becomes zero. The timescale *τ*_*y*_ influences the maximum variance in both cases. **f**. Variance of the activity sensor as a function of the weight-rate alignment timescale, *τ*_*z*_, for different ratios of *τ*_*ρ*_*/τ*_*y*_. For both rapid and slow non-homeostatic plasticity, the activity sensor suppresses fluctuations in the input drive. The maximum variance is achieved for intermediate values of *τ*_*z*_, and its magnitude is defined by the ratio *τ*_*ρ*_*/τ*_*y*_. **g**. The variance of both the regulator and activity sensor at steady state shows an inverse relationship to the number of input synapses *N*.

In addition to heterosynaptic and homeostatic plasticity, individual synapses exhibit various forms of activity-dependent and stochastic plasticity dynamics [22, 38, 91]. Yet, synaptic weight distributions remain largely stable across time under baseline conditions, despite individual variability [64]. Similar stability at a distribution level has been observed for neuronal firing rates [41, 89]. We therefore assume here that individual synaptic plasticity dynamics reshuffle the assignment of weights to rates, without altering their marginal distributions. This formulation avoids assumptions about the nature of such dynamics and allows treating them as a stochastic process.

Based on these assumptions, we define the normalised synaptic weights 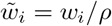 with *ρ* = Σ _*j*_ *w*_*j*_, and express the homeostatic model as:

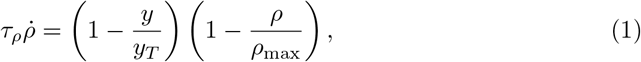

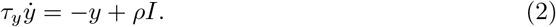

Here, Eq. 1 describes the dynamics of the homeostatic regulator, adjusting the total resource pool, *ρ*, with a timescale, *τ*_*ρ*_. The first term on the right-hand side measures deviations of the activity sensor of the neuron, *y*, from the homeostatic activity target *y*_*T*_. The second term enforces an upper limit on the resource pool equal to *ρ*_max_, motivated by experimental evidence of transcription-dependent restraining of the homeostatic response [105]. The activity sensor (Eq. 2) is a low-pass filtered version of the input drive, 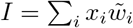, measured in Hz, scaled by the resource pool, *ρ*, over a timescale *τ*_*y*_. The product *ρI* represents the total input drive.

Because both *x*_*i*_ and 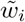 are lognormally distributed [45, 64, 69, 82], the input drive can be approximated by another lognormal distribution, such that 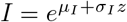 [28]. We can then take the input drive dynamics to be stochastic, by modeling the variable *z* ~𝒩 (0, 1) as an Ornstein–Uhlenbeck process with timescale *τ*_*z*_:

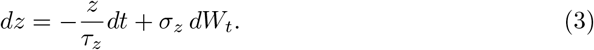

The above equation represents ongoing weight–rate assignment fluctuations. Full derivations and justifications of approximations are provided in the Methods.

### Homeostatic equilibrium is asymptotically stable

First, we evaluate analytically the ability of the model to homeostatically regulate the activity of the neuron. For 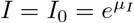, and *z* = 0, we find the two equilibrium points, *h* (homeostatic) and *m* (marginal). Those are associated with reaching the homeostatic target, *y*_*T*_, and the maximum resource pool, *ρ*_max_, respectively:

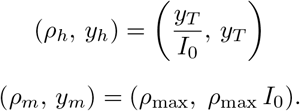

It follows that the homeostatic target can only be reached for sufficiently large maximum total input drive:

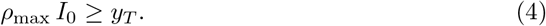

To understand the behaviour of the system, we linearise it by introducing perturbations around the homeostatic equilibrium point *h*. We find that as long as the homeostatic equilibrium can be reached with the maximum total input drive (condition in Eq. 4), the fixed point is stable and the system will always converge towards it. If, on the other hand, the condition is not met, the system settles on the saturation point *m*, which is a saddle. This implies that the saturation point can be reached under strong or sustained perturbations. However, it is unstable, and trajectories can escape it, making homeostasis possible. We conclude that, in line with previous work [99], when synaptic scaling acts at the neuronal rather than the synaptic level, it can prevent runaway dynamics and ensure firing rate homeostasis.

We wondered about the nature of convergence towards the homeostatic equilibrium, and identified two linearly separable dynamic regimes (Fig. 1, b) based on the condition:

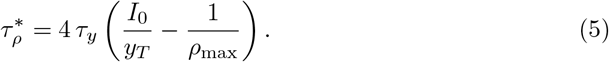

This critical value of the homeostatic regulator, 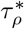, determines the behaviour of the system. Damped oscillations occur when 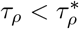 (Fig. 1, c, top), and monotonic convergence for 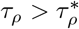 (Fig. 1, c, bottom). In the special case where 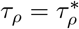, the behaviour is critically damped, with the fastest non-oscillatory return. Our results are summarised in Table 1, and example responses of the neuron are visualised in Fig. 1. d. Therefore, the model can bring its activity to a homeostatic equilibrium, provided it can be reached, and the nature of this convergence is determined by the relationship between the timescales of the activity sensor and the homeostatic regulator. This introduces the first experimentally testable prediction:

**Table 1.** Stability analysis summary.

| Condition | Oscillatory Behaviour | Stability | Notation |
| --- | --- | --- | --- |
| **Homeostatic equilibrium, $h$** |  |  |  |
| $\tau_\rho > 4\tau_y \left( \frac{I_0}{y_T} - \frac{1}{\rho_{\max}} \right)$ | none | stable | sink |
| $\tau_\rho < 4\tau_y \left( \frac{I_0}{y_T} - \frac{1}{\rho_{\max}} \right)$ | damped | stable | spiral sink |
| **Marginal equilibrium, $m$** |  |  |  |
| $y_T \geq I_0 \rho_{\max}$ | none | stable | sink |
| $y_T < I_0 \rho_{\max}$ | none | unstable | saddle |

#### Prediction 1

Assuming that biological neurons operate near the critically damped state, then, manipulating the timescale of the activity sensor by inhibiting or accelerating Ca^2+^ kinetics can push the neuron towards either the monotonic or oscillatory regime. For example, activity blockade drives miniature post-synaptic current (mEPSC) amplitude upward through synaptic upscaling [47, 103]. Upon blocker washout, activity is restored, and the neuron returns to its homeostatic equilibrium. The model predicts that this return should be non-monotonic with mEPSC amplitude transiently overshooting before settling to the baseline when intracellular Ca^2+^ signalling is sufficiently slowed. Conversely, if Ca^2+^ signalling is accelerated, the return should be monotonic.

### Stationary covariance matrix

Ongoing stochastic weight-rate reassignment continuously perturbs the input drive, *I*, causing fluctuations around the homeostatic equilibrium. We wondered how sensitive the system is to such plasticity-driven fluctuations. To that end, we compute the stationary covariance matrix of the linearised system around *h*, reflecting the steady-state variability and statistical dependencies of *ρ, y*, and *z*, in response to ongoing stochastic weight-rate reassignment. We find the structure of the covariance matrix to be:

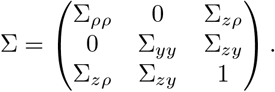

The full form and derivation are provided in the Methods.

Noise enters the system only from *z*, propagating via a clear hierarchy from *z* → *y* → *ρ*. Therefore, the homeostatic regulator only sees a filtered version of the input drive noise, which prevents the resource pool from being destabilised by rapid, transient input fluctuations. Notably, despite the dynamic coupling, the activity sensor and homeostatic regulator are decorrelated at the steady state Σ_*yρ*_ = 0. This suggests that fluctuations in the activity sensor provide no information about the concurrent state of the homeostatic regulator.

As the timescale of the homeostatic regulator increases, *τ*_*ρ*_ → ∞, the variance of the activity sensor, Σ_*yy*_, saturates (Fig. 1, e, top), while that of the regulator, Σ_*ρρ*_, vanishes (Fig. 1, e, bottom). This implies that the regulator becomes unresponsive to fluctuations in the activity sensor. On the other hand, for more rapid homeostatic regulation, *τ*_*ρ*_ → 0, the inverse relationship holds, which reveals a trade-off between the activity sensor and regulator. A slower regulator limits fluctuations in the resource pool, but it allows larger activity variation. Conversely, a faster regulator suppresses activity fluctuations but introduces fluctuations in the resource pool. Biologically, the experimentally observed slow timescale of synaptic scaling sits at the regime where Σ_*ρρ*_ is small, consistent with the observed stability of synaptic weight distributions over time [64].

Furthermore, for very rapid or very slow synaptic plasticity dynamics, *τ*_*z*_ → 0 or *τ*_*z*_ → ∞, the variance Σ_*yy*_ approaches zero, reflecting only a small variation (Fig. 1, f). This reveals that the activity sensor acts as a band-pass filter of synaptic plasticity-driven fluctuations. When such fluctuations are fast, *τ*_*z*_ → 0, the activity sensor only captures the mean input and rapid fluctuations are cancelled out. For very slow synaptic plasticity dynamics, the input drive barely changes, so fluctuations are small. Only at intermediate timescales does the activity sensor react maximally to input noise. Maximising Σ_*yy*_ with respect to *τ*_*z*_, by setting its derivative to zero, we find that the maximum variance of the activity sensor at the homeostatic equilibrium scales linearly with the variance of the input drive, 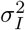. This variance is a function of the standard deviation of both input rates, *σ*_*x*_, and synaptic weights, *σ*_*w*_ (Fig. **??**), suggesting that at the steady-state, fluctuations are primarily driven by input heterogeneity. Therefore, independently of the timescale of synaptic plasticity dynamics, a neuron receiving more homogeneous inputs will show less variability on its activity sensor than one receiving heterogeneous inputs.

Lastly, we note that all terms in the covariance matrix are proportional to *σ*_*I*_. Since 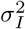 is inversely proportional to the number of synaptic inputs, for larger *N*, fluctuations become smaller (Fig. 1, g). Taking the insights from the covariance matrix together, we can make another testable prediction:

#### Prediction 2

In cultures at homeostatic equilibrium, somatic Ca^2+^ imaging should reveal that neurons with a higher total synapse count show lower variability in somatic Ca^2+^ fluorescence, compared to neurons with fewer synapses. Moreover, neurons with more homogeneous input rates or synaptic weights should show smaller fluorescence variability.

### Parameter fitting under activity blockade reveals a timescale coupling

The activity sensor and homeostatic regulator are each associated with a time constant. Those determine how quickly the neuron can track changes in its own activity and adjust the number of synaptic resources, respectively. We fit these parameters to data from three in-vitro experiments [47, 103] where activity blockade is induced in cultures of visual cortical neurons using TTX. We motivate our choice of the experiments based on the following:

1. We assume that the controlled in-vitro environment of the chosen studies partially isolates synaptic scaling from other homeostatic mechanisms, such as changes in the levels of inhibition [18, 61], intrinsic plasticity [23, 57] and metaplasticity [2, 52, 54, 70]. Additionally, the absence of overall neuronal activity, in contrast to a reduction in activity, as is the case in sensory deprivation, potentially minimises activity-dependent modifications.
2. They provide multiple measurement time points, allowing for higher resolution of the synaptic scaling time course.
3. All three manipulations involve TTX application in comparable experimental settings, namely, bath TTX application [103], and bath and local somatic TTX application in the presence of astrocytes [47], and some overlap between responses is expected.

We find that multiple combinations of the two timescales, *τ*_*ρ*_, *τ*_*y*_, can explain the data (Fig 2, a). Notably, we observe variations between the timescales corresponding to each experimental condition. Our model reproduces the experimental observation that prolonged TTX application to the whole culture with astrocytes (Fig 2, a, middle panel), induces a slower and weaker homeostatic response compared to somatic activity blockade of a single neuron (Fig 2, a, top panel). In the absence of astrocytes, and for sufficiently fast homeostatic response, the timescales of the activity sensor of the neuron can be in the order of hundreds of hours, while slower response requires a much faster reaction from the neuron (Fig 2, a, bottom panel).

**Figure 2.**
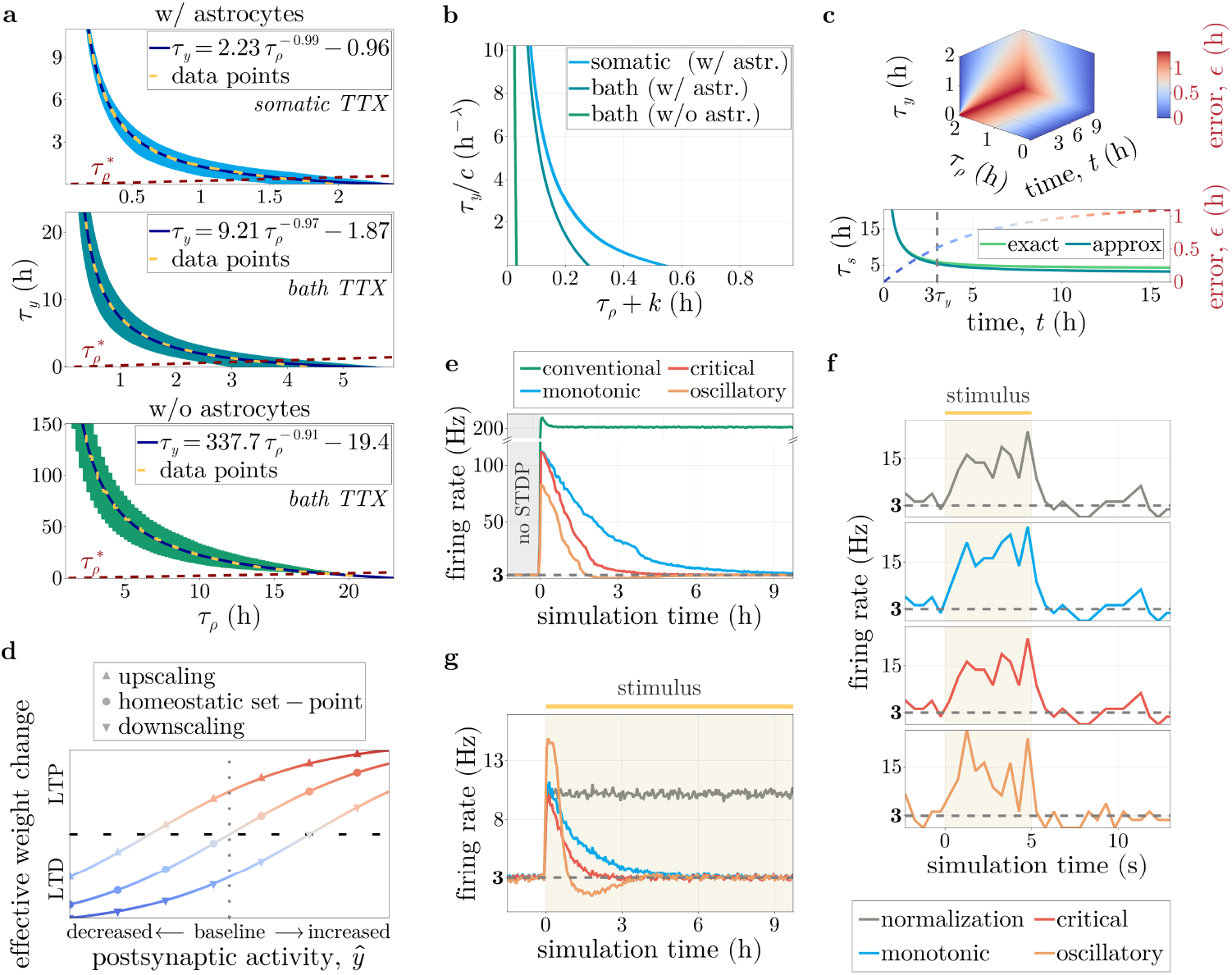
**a.** Timescales of the effector, *τ*_*ρ*_, and activity sensor, *τ*_*y*_, that can replicate the experimental data. Orange dashed lines indicate the critical value for different dynamic regimes. Values below that line indicate monotonic convergence, while those above indicate damped oscillations. The data for *somatic and bath TTX* in cultures with astrocytes (top) are taken from [47] while data for *bath TTX* without astrocytes (bottom) are taken from [103]. **b**. Data collapse for the datasets in **a**. When scaled, the data points do not collapse onto a single curve, suggesting no universal law governing all three experiments. **c**. Accuracy of the approximation of the timescale *τ*_*s*_, around *t* = 0 as a function of time and the two timescales, *τ*_*ρ*_ *& τ*_*y*_ (top). The panel depicts three 2D slices for *τ*_*ρ*_ = 2, *τ*_*y*_ = 0, and *t* = 9. An example case where *τ*_*ρ*_ = 2 and *τ*_*y*_ = 1 (bottom). For relatively small values of the timescales compared to time, *t*, the approximation matches the true value of *τ*_*s*_ with a small error. **d**. Link to metaplasticity. During homeostatic regulation, the effective weight change for a synapse is altered. During synaptic upscaling (downscaling), synapses exhibit a positive (negative) drift, indicating a shift in the postsynaptic activity threshold for potentiation and depression. **e**. Regulation under additive Hebbian plasticity across dynamic regimes. STDP is activated at *t* = 0. Within our framework, convergence is monotonic in the stable regime (blue), fastest at the critical boundary (red), and characterised by damped oscillations in the oscillatory regime (orange). A conventional synapse-specific scaling model (green) fails to prevent runaway activity. Spike trains are binned at 30 min. **f**. Neuronal response to brief 5 sec stimulation. As in a simple normalisation scheme, all regimes produce similar responses, with only minor differences. Here, neuronal spike trains are binned at 500 ms. **g**. Neuronal response to sustained stimulation. Unlike a simple normalisation scheme (grey), our framework compensates for persistent input changes by gradually restoring postsynaptic activity to the homeostatic target of 3 Hz. Spike trains are binned at 30 min.

We wanted to quantify this inverse relationship between the timescales. To that end, we fitted power law functions and found good fits for all three datasets (Fig. 2, a). In particular, we found the function describing the relation between the timescales of the regulator and activity sensor to be

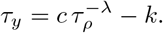

The value of the exponent in all cases is just below one (*λ* > 0.9), suggesting a slightly sublinear coupling. In the following, we assume that *λ* = 1, as an approximation, in an attempt to analyse this coupling, and take

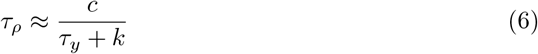

Because the above relationship held across all three cases, we wondered whether the three datasets could collapse into a single curve upon rescaling However, we found no universal law that can describe all three datasets (Fig. 2, b). Nonetheless, we can gain some useful insights. First of all, the offset, *k*, in the denominator introduces a non-linearity for values of *τ*_*y*_ ≪ *k*. Even if we assume that the activity sensor is instantaneous, *τ*_*y*_ → 0, a baseline delay, *k*, is present, reflecting a latency in the upregulation of synaptic resources. That could be delays in the homeostatic regulator of the neuron, for example, due to transcription or receptor trafficking dynamics, or some “buffering” in the activity sensor like calcium/calmodulin-dependent protein kinases (CaMKs) or Ca^2+^ kinetics. Experimental estimates suggest that the delay between transcription factor activation (e.g. Ca^2+^ level changes) and the appearance of mature mRNA is about 10-20 minutes on average [58]. This leads to the third prediction of our model:

#### Prediction 3

Pharmacologically changing the dynamics of CaMKs or Ca^2+^ should change this lag. For example, TTX-induced blockade in combination with CaMKII inhibition should result in a delayed homeostatic response compared to TTX alone. Then, the mEPSC amplitude measured at early timepoints should be indistinguishable from baseline in the former case, while a detectable increase should be observed in TTX-only controls.

To understand why multiple parameter combinations can fit the data, we mimic TTX-induced activity blockade in our model. We assume the neuron is initially at homeostatic equilibrium, with *y*(0) = *y*_*T*_ and *ρ*(0) = *ρ*_0_, and we set the input drive of the neuron in Eq. 2 to zero, *I* = 0. We can then derive the analytical solution for this particular case:

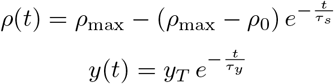

The solution implies that the activity sensor, *y*, decays from the initial homeostatic value, *y*_*T*_, towards zero with a timescale *τ*_*y*_, while the resource pool, *ρ*, increases towards *ρ*_max_ with a time-dependent timescale

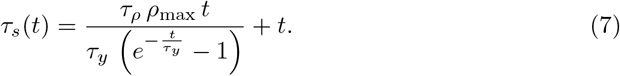

The above equation describes the effective scaling timescale observable in TTX blockade experiments. To get a better intuition about how it relates to the time constants of the activity sensor and regulator, we approximate *τ*_*s*_(*t*) with an asymptotic expansion around *t* = 0:

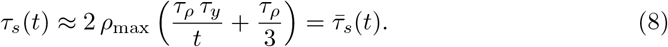

We evaluate its accuracy by defining the error through time, *ϵ*, as the difference between the exact value and our approximation, 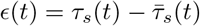. For small *t* relative to the two time constants, the approximation is almost identical to *τ*_*s*_, and progressively diverges as the ratio *τ*_*ρ*_*/τ*_*y*_ increases (Fig. 2, c). This can be explained by the first term in the parenthesis in Eq. 8, which includes the product of the underlying timescales, *τ*_*ρ*_ and *τ*_*y*_, and an inverse proportionality with time, *t*. The second term depends solely on the timescale of the regulator, *τ*_*ρ*_. For *t <* 3*τ*_*y*_, the first term contributes more to the value of *τ*_*s*_, suggesting a strong dependence on the product *τ*_*ρ*_ *τ*_*y*_. For *t >* 3*τ*_*y*_, that value depends more on *τ*_*ρ*_ alone. We illustrate how the error evolves by arbitrarily considering the case for *τ*_*ρ*_ = 2 and *τ*_*y*_ = 1 (Fig. 2, c, bottom). Thus, we identify the dependence of the experimentally observed timescale for synaptic scaling, *τ*_*s*_, on the two underlying timescales of the regulator and activity sensor, in the case of activity blockade using TTX. Taking these into consideration, we can make another testable prediction:

#### Prediction 4

The early phase of TTX-induced synaptic upscaling is governed by the product of the timescales of the activity sensor or the homeostatic regulator, *τ*_*ρ*_*τ*_*y*_. Manipulating either timescale by the same factor should produce an equivalent shift in the observed scaling timescale. For example, inhibiting protein synthesis or slowing Ca^2+^ kinetics by comparable magnitudes should produce indistinguishable effects on mEPSC amplitude at the early timepoints of the homeostatic response (1 − 4 h post-TTX).

### Relation to metaplasticity

Homeostatic synaptic plasticity has increasingly been linked to the concept of metaplasticity [14, 52, 60]. Metaplasticity refers to activity-dependent modification of the synaptic threshold for potentiation and depression [2, 54, 70]. Here, we attempt to make such a connection with our model, drawing on the BCM theory, which utilises a sliding threshold, *θ*, for potentiation versus depression depending on recent postsynaptic activity [13, 120]. We link our mechanism to the notion of metaplasticity through activity-dependent Hebbian synaptic dynamics that include a threshold for potentiation and depression:

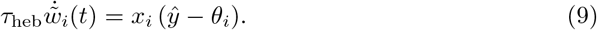

In line with the BCM theory, Hebbian modifications depend on the instantaneous post-synaptic activity, *ý*, which we take to be equal to the total input drive, *ρI*.

However, rather than using a sliding threshold, here we take the threshold, *θ*_*i*_, to be fixed for each synapse. Again, we assume that weights are normalised due to heterosynaptic competition at timescales similar to *τ*_heb_. Then the weight change of a synapse, *i*, is given by 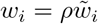 and the weight change is:

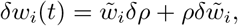

with δ*ρ* the relative change of the total resource pool (Eq. 1), and 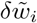 is the relative change of the normalised synaptic weight (Eq. 9). Therefore, the raw weight change of a synapse depends both on rapid, activity-dependent dynamics and slow homeostatic regulation. As a result, our model predicts that during homeostatic upscaling, the observed magnitude of potentiation will be larger compared to when the neuron is at the homeostatic equilibrium, *y* = *y*_*T*_. Similarly, during downscaling, the observed weight change will be more prone to depression. This behaviour resembles that of a sliding threshold, despite *θ*_*i*_ being fixed in our case (Fig. 2, d). Conceptually, increasing the pool of available resources, as is the case during homeostatic upscaling, would allow synapses to be less constrained by competition [100] and be able to acquire more resources during Hebbian potentiation. This leads to another prediction:

#### Prediction 5

Homeostatic regulation of the synaptic resource pool should alter Hebbian learning dynamics. Specifically, during upscaling, the expanded resource pool should lower the threshold for long-term potentiation, such that synapses exhibit larger potentiation magnitudes and weaker stimulation should be required compared to baseline conditions. Conversely, during downscaling, the contracted resource pool should raise the potentiation threshold, favouring depression. Indeed, some experimental evidence is consistent with this prediction, reporting enhanced potentiation during prolonged activity blockade [6, 42].

### Comparison to alternative models

Having established the analytical framework, we now turn to numerical calculations and compare our model with conventional compensatory models of either synaptic normalisation (Eq. 10, introduced below) or synapse-specific synaptic scaling (Eq. 30). We simulate a single leaky integrate-and-fire neuron with excitatory and inhibitory synaptic inputs and log-normally distributed presynaptic firing rates [82]. We set the mean and variance of synaptic weight distributions to match those measured in the cortex [45]. Excitatory synapses are plastic and exhibit Hebbian dynamics in the form of additive spike-timing-dependent plasticity (STDP) [11] (Eq. 24). Inhibitory synapses are static. We implement heterosynaptic plasticity in our numerical simulations as multiplicative normalisation that preserves the sum of synaptic weights equal to a target sum, *W*_*T*_ :

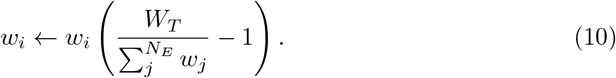

We apply this every 10 ms. The target sum, here, corresponds to the pool of synaptic resources, *ρ*, from our analytical model, and evolves with an effector time constant, *τ*_*ρ*_, as a function of an activity sensor with a timescale *τ*_*y*_. We assume a large enough value for the saturation point of the target sum, and set the homeostatic target rate of the postsynaptic neuron to be equal to the mean of the distribution of input firing rates. Then, the critical boundary (Eq. 5) is approximately:

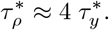

Using the above relationship, we choose three sets of values for the two timescales, making sure that both are sufficiently larger than those associated with STDP (Table 2). The three sets correspond to the two dynamic regimes, namely the monotonic and oscillatory, and the critical boundary between them.

**Table 2.** Synaptic scaling parameters for each case.

| time constant | oscillatory | critical | monotonic | conventional |
| --- | --- | --- | --- | --- |
| regulator ($\tau_\rho$) | 30 min | 1 h | 2 h | 1 h |
| activity sensor ($\tau_y$) | 30 min | 15 min | 10 min | 15 min |

We first compare the behaviour of the postsynaptic neuron operating using our model, with that of a synapse-specific synaptic scaling that acts on individual synapses (Fig. 2, e). After an initial calibration period without plasticity, we switch on the STDP. Activity-dependent modifications lead to the potentiation of some input synapses and eventually to an increase in the activity of the neuron. This triggers the homeostatic response, regulating the postsynaptic firing rate to the target set-point. As predicted by the analytical model, we observe differences in the behaviour of the neuron under the different regimes. In the monotonic regime and at the critical boundary, the homeostatic response brings the firing rate of the neuron steadily back to its target rate. In the oscillatory regime, we observe the expected damped oscillations around the target rate. Regardless of which regime the neuron operates in, we observe that the firing rate returns to its homeostatic set-point, counteracting instabilities that arise from Hebbian plasticity.

In contrast, we find that the synapse-specific model acting on biological timescales cannot compensate for the runaway growth due to Hebbian dynamics, in line with previous work [124]. The two forces acting on the synapse differ substantially in magnitude, and the slow, negative homeostatic feedback is too weak to balance the positive Hebbian drive. Compensation occurs only when the homeostatic force becomes comparable in magnitude to the Hebbian one. By that point, however, the system has already drifted and settled far from the homeostatic fixed point. To address this, some models [106, 122] utilised an integral controller that accumulates the error over time, bringing neuronal activity back to its target firing rate. As previously reported [106, 121], we found that incorporating such a mechanism can suppress certain oscillations. However, it still requires parameter fine-tuning and does not abolish the requirement for similar orders of magnitude between the STDP and synaptic scaling timescales [121]. Thus, we illustrate numerically that when homeostatic forces act at the neuron-level, the system can effectively compensate for the rapid positive-feedback dynamics without requiring tight timescale matching.

We next compare our model with multiplicative normalisation alone. Normalisation enforces a constraint on the total synaptic weight, but is still sensitive to the presynaptic activity, which should allow transient responses to varying inputs. We simulate the same LIF neuron and induce stimulation through an increase in the firing rates of 250 random input neurons. For brief stimulation periods of 5 s, the neuronal responses are quite similar across all cases, suggesting almost no differences between the two models (Fig. 2, f). However, when stimulation is sustained over many hours (Fig. 2, g), normalisation is unsuited to regulate the activity, and the neuron remains sensitive to the increased frequency of the input. By contrast, with our model, the synaptic weights are scaled down to bring the postsynaptic firing rate to its homeostatic set point. Again, the main difference between the dynamic regimes is the nature of convergence towards equilibrium. We show that combining synaptic normalisation and homeostatic synaptic scaling within a single framework enables rapid responses to stimulation at short timescales and regulation of activity at much longer timescales, while preventing pathological activity.

### Synaptic competition under local resource constraints produces more realistic weight distributions

Although synaptic competition through normalisation enforces a constraint on the sum of synaptic efficacies, it cannot stabilise synaptic distributions. In particular, nothing prevents individual synapses from monopolising the total synaptic strength, leading to bimodal distributions [106] (Fig. **??**). Synaptic competition and conservation of total synaptic strength have been reported among neighbouring synapses, typically within the same dendritic branch [7, 15, 17, 20, 27, 75, 98, 112]. Such within-branch competition can potentially have computational benefits, by supporting multiple feature representations in a single neuron at separate dendritic domains [51]. We wondered whether such local restrictions influence the homeostatic response and the resulting synaptic distributions. We explore this by introducing spatial constraints such that only nearby synapses compete for shared resources. Specifically, we partitioned synapses into *N*_*c*_ ≥ 1 dendritic compartments, and updated our normalisation rule from Eq. 10 to:

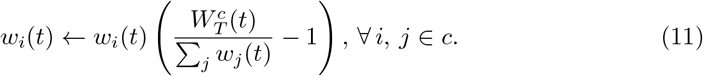

Here, 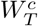 is the normalisation target for all excitatory input weights to the dendritic compartment, *c*. Again, we apply the above normalisation every 10 ms. For simplicity, we assume that synapses within a compartment share a common normalisation target and that each compartment contributes a fixed fraction to the total synaptic strength. We further assume that as individual synapses approach the local target, 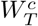, they exhibit stronger competition, limiting potentiation. We implement this by modifying the STDP rule by introducing a soft upper bound on potentiation in addition to weight-dependent depression (Eq. 25). We performed simulations, varying the number of compartments, *N*_*c*_, and compared the results of the modified STDP to those of one that has weight-dependent depression but without an upper bound (Fig. 3, a–c). Increasing the number of compartments progressively constrained the initial STDP-driven activity burst. This follows naturally from the reduced size of the local resource pool, 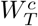, which limits individual synaptic potentiation. Note that with 200 dendritic compartments, for example, each sum is shared by only 5 synapses. Incorporating the upper bound restricted this transient response even further (compare insets in Fig. 3, a). The distribution of synaptic weights also acquires different shapes, depending on the STDP rule. In both cases, the mean weight increased with the number of compartments (Fig. 3, b, solid lines). However, when STDP included both weight-dependent depression and an upper bound, the variance decreased with increasing *N*_*c*_. Without the upper bound, the variance remained approximately constant (Fig. 3, b, dotted lines).

**Figure 3.**
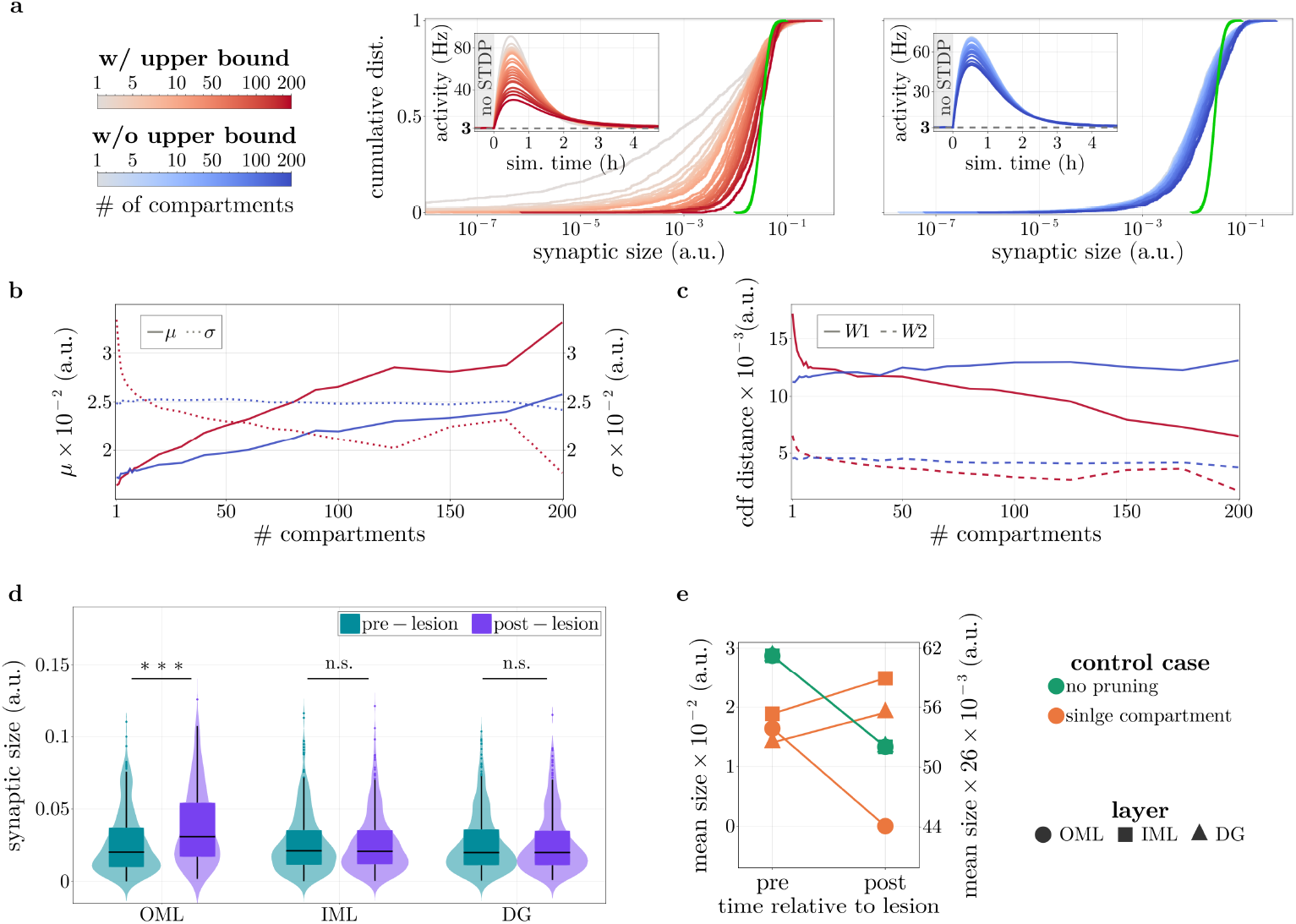
**a.** Comparison of resulting distributions as a function of number of dendritic compartments for STDP with (red) and without (blue) an upper bound, and the corresponding post-synaptic activity traces of the neuron (insets). Increasing the number of compartments progressively (*N*_*c*_ ∈ [1, 2, …, 10, 20, …, 100, 125, …, 200]) attenuates the STDP-induced activity burst in both conditions, with a stronger reduction when an upper bound is imposed. Here, the experimentally observed distribution of synapses of a single neuron (green) is scaled to the mean of the distributions with 200 dendritic compartments for each case. **b**. Mean (left axis, solid lines) and variance (right axis, dotted lines) corresponding to panel a. Without an upper bound (blue), the variance remains approximately constant as the number of compartments increases, but it decreases when STDP includes an upper bound (red). In both cases, the mean synaptic weight increases with compartment number. **c**. Wasserstein (*W* 1, solid lines) and tail-restricted Wasserstein-2 (*W* 2, dashed lines) distances plotted against the number of dendritic compartments. When STDP lacks an upper bound (blue), both metrics remain largely unchanged. In contrast, incorporating an upper bound (red) leads to a progressive reduction in both distances. Here, *W* 2 is computed for the largest 5% of synapses. **d**. Local scaling of synapses results from the redistribution of building blocks. Denervation of one-third of synaptic inputs to the outer molecular layer (OML) produces an increase in the distribution of synaptic sizes post-lesion. This increase is restricted locally as there are no changes in the mean synaptic size in the inner molecular layer (IML) or the dentate gyrus (DG); Mann-Whitney test, p-values: 7.8*e*^−8^ (OML), 0.9 (IML), 0.96 (DG). **e**. Control cases for local synaptic scaling. In the case of a single compartment (left axis, orange), the mean synaptic size decreased in OML, which was compensated by an increase in DG and IML. When synaptic pruning is omitted (right axis, green), a slight uniform reduction is observed across all layers.

To assess how well these distributions matched experimental observations, we sampled a log-normal distribution with mean and variance reported in [64] (Fig. 3, a, green curves). These sampled distributions were rescaled to match the same mean as the simulation results while preserving their coefficient of variation. We then quantified discrepancies using the Wasserstein distance and, to probe differences in the upper tail, the tail-restricted Wasserstein-2 distance computed on the top 5% of synapses. Incorporating the upper bound reduced both distance measures, while without it the distances remained largely unchanged across compartment numbers (Fig. 3, c). However, we find that our simplified STDP rule cannot fully capture the complexity of biological synaptic dynamics. Nonetheless, these results suggest that locally restricted redistribution among a few synapses, combined with stronger competition near resource limits, yields distributions that more closely resemble empirical data, and predict that synapses approaching their local resource ceiling should exhibit reduced Hebbian potentiation.

### Local redistribution accounts for experimentally observed local scaling

Homeostatic synaptic scaling is often considered to be a global mechanism affecting all synapses of a neuron [102, 103, 112]. However, experimental evidence suggests that this might not be the case. For example, granule cells situated in the dentate gyrus (DG) have their proximal dendrites predominantly in the inner molecular layer (IML) and their distal dendrites in the outer molecular layer (OML) of the DG. Eliminating entorhinal inputs that terminate in the OML results in synaptic loss on the distal dendrites, while synapses found on proximal dendrites remain unimpaired [108]. Thus, the selective removal of synaptic inputs targeting the distal dendrites of granule cells induces layer-specific homeostatic upscaling of excitatory synapses in OML. We wondered whether our model could account for this surprising observation.

We mimic the experiment on local synaptic scaling [108], by dividing synapses in our model neuron into *N*_*c*_ = 100 dendritic compartments, and assigning each compartment to a specific layer, namely DG, IML and OML. We then simulate the neuron with STDP until it reaches equilibrium and selectively silence one-third of synaptic inputs in the OML layer, thereby replicating entorhinal denervation. As expected, we observed that those synapses were rapidly reduced to the low hard bound due to Hebbian depression. Since synapses that exhibit long-term depression are eliminated unless this is compensated by presynaptic activity [16], we randomly pruned silenced synapses (one every 500 ms arbitrarily), by setting their weight to zero. Synapses still receiving presynaptic input are potentiated due to STDP and normalisation, reshaping the synaptic weight distribution in the OML. The resulting distribution is analogous to that predicted by local synaptic scaling (Fig. 3, d), though with fewer synapses that are larger in size. Thus, layer-specific synaptic scaling emerges naturally within our framework as a consequence of local resource redistribution, while homeostatic regulation remains a cell-wide process.

To understand which aspects of the model are responsible for the observed scaling effect, we run two control simulations. First, we begin by omitting synaptic pruning. We observe a mild uniform reduction in the mean sizes across layers (Fig. 3, e, green). This results from the slightly increased activity after the lesion, due to potentiation of the intact OML synapses. Second, we assess the importance of dendritic compartmentalisation by switching back to neuron-wide normalisation. In this case, and in direct contrast to the experimentally observed dynamics, the mean synaptic size decreased in OML, while in IML and DG it increased (Fig. 3, e, orange). These results indicate that local synaptic scaling arises from redistribution of synaptic resources rather than uniform global scaling, and requires both compartmentalisation and pruning. Previous studies have already established a correlation between within-branch spine loss and an increase in the size of the remaining spines [7]. Our results suggest that the timescale with which the local scaling effect occurs is not related to conventional homeostatic synaptic scaling but rather to the redistribution of synaptic resources after synaptic death. Consistent with our interpretation, the timescales of synaptic size increase after entorhinal denervation match those of the reduction in synaptic spine density post-lesion [108, 109]. This interpretation yields another testable prediction:

#### Prediction 6

If layer-specific synaptic scaling reflects local resource redistribution driven by synapse elimination, then its timescale should be governed by the rate of spine elimination rather than by the kinetics of the activity sensing or protein synthesis, predicting a dissociation between the two processes. Pharmacological manipulation of somatic Ca^2+^ signalling or protein synthesis should leave the timescale of post-denervation synaptic size increase largely unaffected. Interventions that accelerate or delay spine elimination should shift the timescale of the size increase proportionally.

## Discussion

We have presented a simple unified framework for synaptic scaling derived from the idea of a limited synaptic resource pool. We suggest that firing rate homeostasis is established at two timescales, in line with previous perspectives [123]. At fast timescales, heterosynaptic competition rapidly redistributes synaptic resources, counterbalancing fluctuations in input drive arising from Hebbian plasticity. This competition arises naturally from the limited availability of shared resources [100]. On a slower timescale, persistent deviations from a firing rate set-point regulate the total resource pool, ensuring convergence to homeostatic equilibrium. The homeostatic response requires the neuron to sense its own activity and adjust its synaptic weights accordingly. In biological neurons, certain calcium/calmodulin-dependent protein kinases (CaMKs) can act as an activity sensor, via measuring intracellular Ca^2+^ levels [47, 97, 101]. Changes in CaMK activity have indeed been linked to homeostatic responses [34, 49, 94], and polarity proteins may encode the sign of the response by being differentially regulated during upscaling and downscaling [24]. The downstream regulatory response appears to be transcription-dependent, affecting the synthesis, turnover, and degradation of synaptic proteins [46, 47, 84, 85, 90, 92, 96]. Such processes modulate the size of the available resource pool. Deliberately avoiding strong assumptions about the molecular identity of the activity sensor or regulator, our framework instead characterises their dynamic relationship, offering insights into the stability of synaptic scaling and generating experimentally testable predictions.

Unlike alternative models, which require a rapid homeostatic response matching the timescale of Hebbian plasticity, our framework guarantees convergence regardless of the difference in magnitude between these timescales. Consistent with previous work highlighting the need for cell-wide rather than synapse-specific regulation [99], this architecture permits transient, information-carrying fluctuations in firing rate without engaging the homeostatic regulator — only sustained deviations from the set-point trigger a regulatory response. Moreover, our analysis suggests the existence of two dynamic regimes for synaptic scaling. Based on the relative ratio of the two underlying timescales of the activity sensor and homeostatic regulator, convergence to equilibrium is either monotonic or oscillatory. Which regime cortical neurons operate in remains unclear from existing data. Chronic pharmacological activity blockade in vitro is typically not accompanied by restoration of firing rates, precluding direct observation of the return to equilibrium. In vivo evidence, by contrast, points to damped oscillations during homeostatic regulation [41, 78, 118]. Whether these reflect synaptic scaling specifically or arise from other compensatory, Hebbian, or stochastic processes remains unresolved. Determining which regime operates in cortical neurons will require targeted experiments. The first prediction of our model addresses this directly. Altering the ratio of the two underlying timescales during TTX-induced activity blockade, for example, by pharmacologically manipulating Ca^2+^ kinetics, should shift the nature of the homeostatic response upon TTX washout from monotonic to oscillatory or vice versa.

Furthermore, we find that, given rapid heterosynaptic competition, input fluctuations do not destabilise the system. The activity sensor can filter rapid fluctuations at the homeostatic equilibrium, preventing transient perturbations from propagating to the homeostatic regulator. This provides a principled explanation for why homeostatic regulation operates on much slower timescales than Hebbian plasticity in biological neurons. Rapid Hebbian plasticity allows the system to adapt quickly to new experiences, while competition counteracts rapid runaway synaptic dynamics. The slow homeostatic regulator remains insulated from transient fluctuations, preventing extreme deviations in cortical activity without interfering with learning. At equilibrium, the magnitude of residual fluctuations in both the activity sensor and the regulator is primarily determined by input heterogeneity, regardless of the timescale of Hebbian plasticity. Consistent with the law of large numbers, neurons with more synaptic inputs should exhibit smaller fluctuations, yielding a directly testable prediction (Prediction 2).

We find that the experimentally measured timescale of TTX-induced synaptic scaling is the product of two underlying timescales, those of the regulator and activity sensor. The existence of multiple combinations of parameter values that can explain the data reveals a timescale coupling implying two biologically plausible extremes. A rapid but weak response, in which the neuron quickly detects activity loss but slowly adjusts resource production and degradation, or a slow but strong response, in which activity deviations accumulate gradually before triggering a robust regulatory reaction. Disentangling these two modes experimentally would require independent manipulation of the sensing and regulatory timescales, for example, through pharmacological targeting of Ca^2+^ kinetics and protein synthesis, respectively (Predictions 3 and 4). Lastly, a further consequence of our framework is that the size of the synaptic resource pool should gate Hebbian plasticity, lowering the threshold for long-term potentiation during upscaling and raising it during downscaling (Prediction 5).

### Spatial resource constraints and local synaptic scaling

Competitive dynamics over synaptic resources are expected to act preferentially among spatially proximate synapses. This is consistent with experimental evidence showing that total synaptic strength remains approximately constant within dendritic branches despite ongoing spine dynamics [15]. Such branch-specific competition can expand the information processing capabilities of individual neurons by supporting multiple independent representations across dendritic domains [7]. However, the spatial organisation of heterosynaptic interactions remain incompletely understood. Some studies report distance-dependent modifications, with adjacent synapses following the direction of change of the stimulated synapse while more distant synapses change in the opposite direction [59, 98]. Others describe a broader, stimulus-selective organisation, in which synapses that share similar input preferences are co-regulated while those that do not, change in the opposite direction [29, 65]. It might be the case that the regulatory schemes are neuron or brain-area-specific. For instance, input synapses of visual cortical neurons with similar orientation preferences are broadly distributed on the dendritic tree [48], while those of hippocampal neurons remain clustered together, suggesting also distinct functions for each distribution scheme [59]. In our simulations, we assumed that each dendritic compartment contributes a fixed fraction to the total resource pool. This is unlikely to be true. In reality, normalisation subunits likely overlap, with individual synapses participating in more than one competitive pool. Protein synthesis, occurring both at the soma and locally at dendrites in a metabolically regulated manner [9], may further modulate the size of the local resource pool. Nonetheless, since local protein synthesis can be induced at individual dendrites [95], each compartment likely has access only to a spatially restricted resource pool, providing a biophysical basis for the compartmentalisation assumption. If compartment contributions were to drift substantially over time, nothing would in principle prevent one compartment from dominating the total synaptic drive. While our framework cannot capture the full complexity of heterosynaptic spatial organisation, the core principles remain valid. Despite the simplified compartmentalisation assumption, restricting competition to local dendritic domains yields synaptic weight distributions that more closely reflect those measured in cortical neurons. Extending the model to incorporate more heterogeneous competitive architectures is a natural direction for future work.

Synaptic scaling induced locally rather than cell-wide [7, 43, 95, 108] also finds a natural explanation within our framework. Rather than reflecting a locally generated homeostatic response, it emerges from the redistribution of a locally constrained resource pool. Synapse elimination releases resources that are redistributed among remaining synapses, increasing their weights and boosting excitatory drive. Consistent with this interpretation, spine density decreases have been observed before spine size increases following both monocular enucleation [7] and prolonged TTX-induced activity blockade [116], suggesting that surviving spines enlarge because they inherit resources released by eliminated synapses rather than through an independent scaling mechanism. The timescale of this effect is therefore set by the rate of spine elimination rather than by conventional scaling dynamics [116]. As individual synapses grow larger through this process, the local maximum resource pool increases, but the number of synapses that can be simultaneously sustained decreases [116, 117]. If local synaptic scaling reflects resource redistribution following synapse elimination, then changing the kinetics of activity sensing or protein synthesis through pharmacological manipulation of Ca^2+^ signalling or spine elimination rates should not affect the timescale of the homeostatic response (Prediction 6).

### What is the timescale of synaptic scaling?

In our framework, the complex molecular dynamics underlying synaptic scaling are deliberately reduced to two processes, an activity sensor and a homeostatic regulator, each characterised by a single timescale. In reality, however, the homeostatic response is at least biphasic and involves multilayered molecular interactions [84, 85, 94]. What, then, is the timescale of the homeostatic response, and which factors shape it? Experimental measurements typically rely on synaptic receptor fluorescence or mEPSC amplitudes as proxies for synaptic strength [5, 40, 47, 53, 93, 103]. The most direct molecular candidates underlying these measurements are *α*-amino-3-hydroxy-5-methyl-4-isoxazolepropionic acid receptors (AMPARs), the primary mediators of fast excitatory synaptic transmission. Anchored to postsynaptic density (PSD) scaffolding proteins [86], their synaptic abundance correlates tightly with synaptic strength [66, 74, 77]. AMPARs undergo rapid turnover, and their concentration at dendritic spines is modulated by activity-dependent [3], heterosynaptic [4], and homeostatic [115] forms of plasticity. During synaptic scaling, changes in AMPAR abundance involve transcriptional regulation [105], modulation of metabolic half-life [77], local dendritic synthesis [50], and changes in turnover and anchoring rates at PSD sites [92, 96]. However, the number of synaptically anchored receptors also depends on other factors, such as the availability of PSD scaffold proteins [4], which are themselves homeostatically regulated but on different timescales [107]. This suggests that multiple parallel processes impose dynamic constraints on one another, shaping the overall time course of the response. Moreover, during the initial phase of synaptic scaling, changes are predominantly post-translational, involving altered trafficking and anchoring rates [85], while total AMPAR levels change over the course of approximately 24 hours [84, 92]. These multilayered interactions impose dynamic constraints on one another, making precise experimental estimates of the synaptic scaling timescale inherently difficult to obtain.

A further challenge is isolating synaptic scaling from other compensatory processes. Cortical neurons are regulated by multiple homeostatic mechanisms acting across overlapping timescales [25, 31, 61, 94]. Since chronic activity blockade, the most common experimental approach to inducing synaptic scaling [71], triggers many of these processes simultaneously [49], attributing the observed response specifically to synaptic scaling remains challenging. Moreover, the expression and dynamics of synaptic scaling are highly heterogeneous across the cortex. Not all neurons homeostatically regulate their activity [8], and among those that do, the dynamics vary substantially with cell type [35], brain region [33], brain state [41, 78], cortical layer [52], and age [25]. The time course of scaling is further shaped by miniature synaptic transmission [95] and by interactions with local circuits and astrocytes [36, 47, 79]. A recent study additionally reported non-monotonic regulation of synaptic strength following activity deprivation, with signs of oscillatory dynamics [94], consistent with our model’s prediction of an oscillatory regime, and highlighting the need for more temporally resolved tracking of the homeostatic response. Collectively, these factors make characterising the time-dependent evolution of synaptic scaling a considerable experimental and theoretical challenge.

## Methods

### Analytical model

We consider a single neuron that receives input from *N* presynaptic neurons with firing rates, *x*_*i*_, where ln 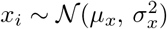. We assume normalized relative synaptic weights, 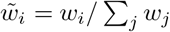 with ln 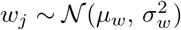. We verify that the normalisation yields approximately lognormal weights, 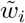 (Fig. **??**). We define the input drive of the neuron to be 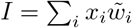. Non-homeostatic plasticity dynamics operating on these normalised weights reshuffle their assignment with input rates. We make no assumptions about how non-homeostatic plasticity influences the assignment of weights to rates and treat pairs of (*x*_*i*_, *w*_*i*_) as i.i.d. Because the product of independent lognormal distributions is also lognormal, and the distribution of the sum of lognormal terms is itself approximately lognormal [28], we take 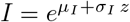, with *z* a Gaussian variable. To find expressions for *µ*_*I*_ and *σ*_*I*_, we first exploit the normalisation of weights, and without loss of generality set 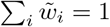. For large enough *N*, the expected value 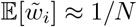 (Fig.**??**), and the log mean shifts to 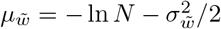. In contrast, the log standard deviation of the weights is preserved, 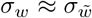 (Fig. **??**). We then combine this with the

Fenton-Wilkinson approximation [28], to find:

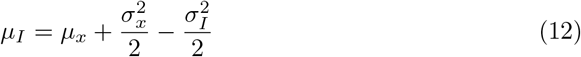

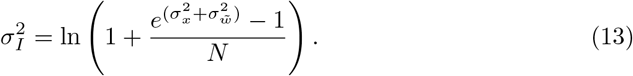

We validate the accuracy of our approximation numerically using a Monte Carlo method and experimentally observed distributions (Fig. **??**). Then, we can write the whole system as:

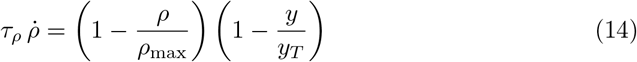

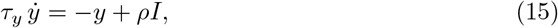

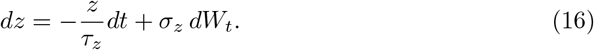

We take the variable *z* to be stochastic, representing ongoing plasticity through weight-rate assignment fluctuations. We model this using an Ornstein-Uhlenbeck process. Here, *τ*_*z*_ represents the timescale of non-homeostatic plasticity dynamics. To ensure that the Ornstein-Uhlenbeck process samples the desired distribution of *I* independently of the choice of *τ*_*z*_, we require the stationary variance of *z* to be equal to one, which gives the constraint 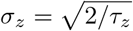.

### Fixed points

We first find the fixed points, for 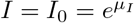, and *z* = 0, by setting 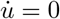 and 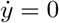:

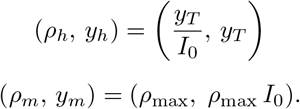

The existence of the homeostatic equilibrium, *h*, is conditioned by:

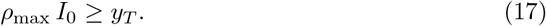

#### Fixed point

*h* We linearise the system by introducing perturbations around the homeostatic fixed point *h*:

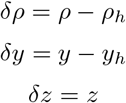

Using a Taylor expansion, we first compute δ*I* = *I*_0_*σ*_*I*_ δ*z*, for small *z* and write the corresponding differential equations:

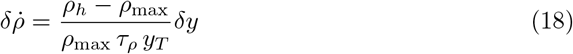

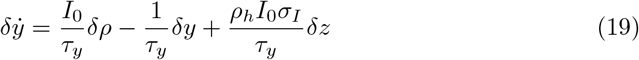

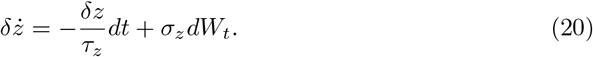

The linearised equations above in matrix form are given by:

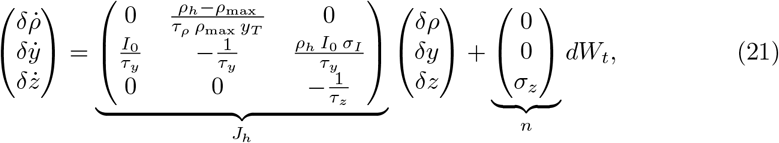

with *J*_*h*_ the Jacobian matrix evaluated at the fixed point *h*, and *n* the noise vector.

### Stability and oscillatory conditions

We observe that the Jacobian, *J*_*h*_, is a block upper-triangular matrix. The first two eigenvalues of the system come from the upper 2 *×* 2 block with characteristic polynomial:

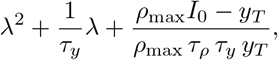

while the third one is 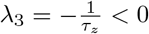. The latter suggests that the stochastic part is always stable. To characterise the system, we first write the 2 *×* 2 Jacobian:

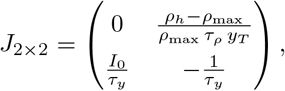

and the corresponding trace and determinant:

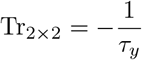

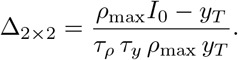

The determinant Δ_2*×*2_ can only be negative when *ρ*_max_ *I*_0_ *< y*_*T*_. This is exactly the condition in Eq 4. Thus, as long as the homeostatic equilibrium can be reached with the maximum total input drive, the fixed point *h* is always stable. To understand the nature of convergence towards equilibrium, we find the condition for which the discriminant, *D*, is equal to zero:

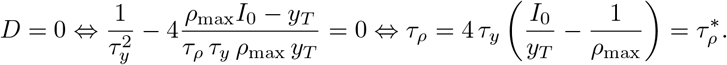

For values of 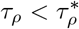, the discriminant is negative, *D <* 0, which gives complex eigenvalues *λ*_1_, *λ*_2_, and the behaviour of the system is characterised by damped oscillations. When 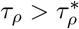, convergence is monotonically stable. In the special case where 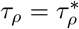, the behaviour is critically damped, with the fastest non-oscillatory return.

#### Stationary covariance matrix Σ

For the linearised system above, we use the algebraic Lyapunov equation [32]:

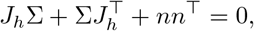

and compute the covariance matrix:

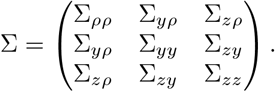

For simplicity, we rewrite the Jacobian as:

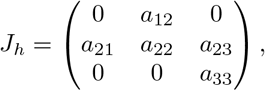

with:

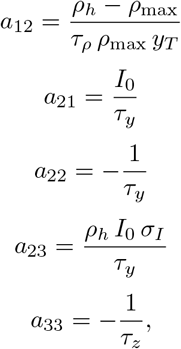

and solve the system of equations below:

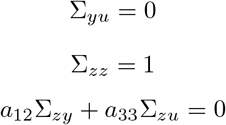

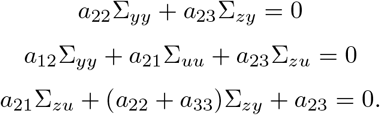

We find the covariance matrix to be:

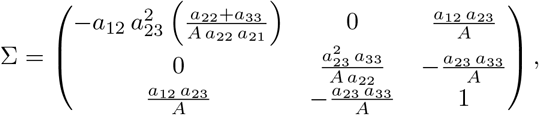

with 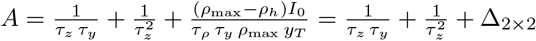.

#### Fixed point *m*

Evaluating the Jacobian at the saturation point, we get:

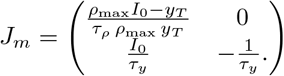

Because *J*_*m*_ is lower triangular, the eigenvalues are given by the diagonal terms, 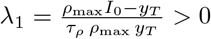 and 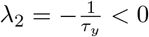. Similarly to the fixed point *h*, the stochastic part decouples with 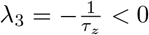. Because the eigenvalues have opposite signs, and the determinant 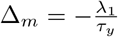, the fixed point *m* is a saddle. This implies that the saturation point *m* can be reached under strong or sustained perturbations. However, in the biologically relevant regime where *ρ*_max_*I* ≥ *y*_*T*_, it is unstable, and thus trajectories can escape it, and homeostasis becomes possible.

### Neuronal dynamics

Membrane dynamics followed the leaky integrate-and-fire (LIF) model given by

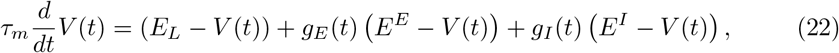

where *V* is the membrane voltage with a time constant *τ*_*m*_, *E*_*L*_ is the resting potential, *g*_*E*_, *g*_*I*_ are the total excitatory and inhibitory input conductances, and *E*^*E*^, *E*^*I*^ are the excitatory and inhibitory reversal potentials, respectively. When the voltage reaches a threshold *V*_*T*_, a spike is triggered and the voltage is reset to *V*_reset_. Following a spike, the neurons exhibit an absolute refractory period, *τ*_ref_, during which they are not able to emit another spike. Parameter values are summarized in Table 3

**Table 3.** Membrane and synaptic dynamics parameters.

| Parameter | Description | Value |
| --- | --- | --- |
| $V_T$ | Voltage threshold | $-50$ mV |
| $E_L$ | Resting potential | $-60$ mV |
| $E^E$ | Reversal potential for E synapses | $0$ mV |
| $E^I$ | Reversal potential for I synapses | $-80$ mV |
| $V_{\text{reset}}$ | Reset voltage | $-60$ mV |
| $\tau_m$ | Membrane time constant | $20$ ms |
| $\tau_{\text{ref}}$ | Absolute refractory period | $2$ ms |
| $\tau_E$ | Excitatory synapses time constant | $2$ ms |
| $\tau_I$ | Inhibitory synapses time constant | $5$ ms |

### Synaptic dynamics

Synaptic conductances were modelled as exponentially decaying variables:

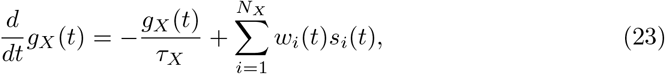

where *X* ∈ (*E, I*), *τ*_*X*_ is the synaptic decay time constant, *w*_*i*_ is the synaptic strength from presynaptic neuron, *i*. The spike train of each neuron *i* is defined as the sum of Dirac delta functions, 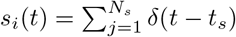, with *t*_*s*_ the spike times for *s* = 1…*N*_*s*_.

### Synaptic plasticity

#### Hebbian plasticity

Excitatory synaptic weights exhibit Hebbian dynamics according to spike-timing-dependent plasticity (STDP). That was either additive

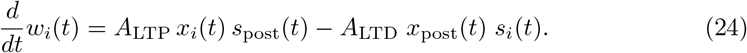

For Fig. 3, we included a soft upper bound and/or weight-dependent depression

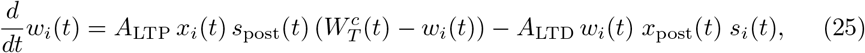

where 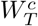 is the target some of dendritic compartment, *c*, the synapse, *i*, belongs to (see next section). *A*_LTP_, *A*_LTD_ are the amplitudes of potentiation and depression, respectively, and *s*(*t*) is as described above. To keep track of previous neuronal activity we use exponentially decaying spike traces *x*(*t*). When a neuron, *i*, spikes, the value of its trace is increased by 1 and decays back to zero with a time constant, *τ*_*H*_ :

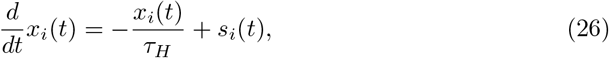

with *H* ∈ (LTP, LTD), denoting long-term potentiation or depression.

### Heterosynaptic plasticity

We group synaptic inputs into *N*_*c*_ ≥ 1 dendritic compartments. We implement heterosynaptic plasticity by normalising synaptic weights over the sum of the compartments they belong to:

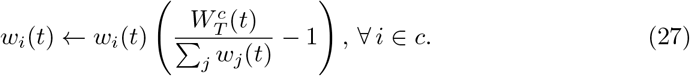

Here, 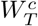 is the normalisation target for all excitatory synapses of compartment *c*, and 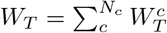. For simplicity, during scaling, we preserve the ratio of 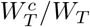 constant. We apply the normalisation every 10 ms.

### Synaptic scaling

Within our framework, the target sum of all synaptic weights is not fixed but given by

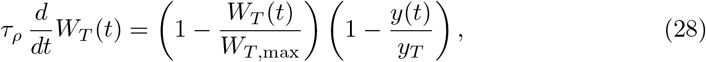

where *τ*_*ρ*_ is the timescale of the homeostatic regulator, *W*_*T*,max_ is a saturation point for the summation target. *y*_*T*_ is the homeostatic target of the postsynaptic for neuron and 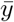 is a running average of its firing rate with time-constant *τ*_*y*_:

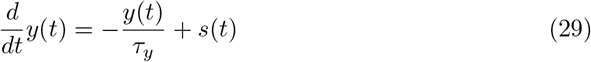

For comparison, we use a conventional synaptic scaling model that multiplicatively adjusts individual synaptic weights as

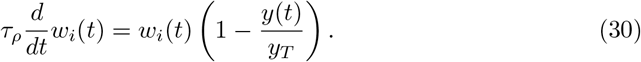

## Simulations

For our numerical calculations, we simulated a single postsynaptic neuron receiving input from *N*_*E*_ excitatory and *N*_*I*_ inhibitory neurons. Incoming spike trains are generated from independent Poisson processes with firing rates drawn from a lognormal distribution with means *µ*_exc_ and *µ*_inh_, and variance of the log, 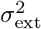. Parameter values are summarised in Table 4. All simulations were implemented in Julia [10].

**Table 4.** Plasticity and input parameters. Parameter values with a ⋆ are taken from [12], and with *†* are from [45].

| Parameter | Description | Value |
| --- | --- | --- |
| $A_{\text{LTP}}\star$ | Amplitude for LTP | 0.0096 |
| $A_{\text{LTD}}\star$ | Amplitude for LTD | 0.0053 |
| $\tau_{\text{LTP}}\star$ | Time constant for LTP | 16.8 ms |
| $\tau_{\text{LTD}}\star$ | Time constant for LTD | 33.7 ms |
| $y_T$ | Homeostatic target rate | 3 Hz |
| $N_E$ | Number of excitatory input neurons | 1000 |
| $N_I$ | Number of inhibitory input neurons | 250 |
| $\mu_{\text{exc}}\dagger$ | Mean rate of excitatory inputs | 3 Hz |
| $\mu_{\text{inh}}$ | Mean rate of inhibitory input | 5 Hz |
| $\sigma_{\text{ext}}^2\dagger$ | Variance of the log of external inputs | 1.04 |

## Code availability

The code is available online at https://github.com/p-vlachos/unified-synaptic-scaling.

## Data acquisition

We acquire the data we fit in Figure 2, a, directly from the published figures in the corresponding publications using the g3data data extraction tool. For bath TTX without astrocytes data were collected from the inset barplot in Figure 2a [103]. For the cases where astrocytes were present, data for bath TTX were acquired from Figure 2A [47], while for somatic TTX we fitted the model to both Figures 3B and S1 [47].

## Acknowledgments

This work was supported by the Deutsche Forschungsgemeinschaft (German Research Foundation, DFG), under Germany’s Excellence Strategy (EXC 3066/1 “The Adaptive Mind”, Project No. 533717223) and SPP 2041 (“The dynamic connectome: dynamics of learning”, Project number 347573108), and the Johanna Quandt foundation (JT).

## Competing interests

The authors declare no competing interests.

## Notes

### Competing Interest Statement

The authors have declared no competing interest.

### Summary of Updates

The manuscript has been substantially revised and extended in several respects. The analytical treatment of the model has been extended to include stochastic Hebbian plasticity, providing a more complete theoretical characterisation of the framework. The steady-state analysis has been made more rigorous through the introduction of a covariance matrix analysis, which formally characterises the variability and statistical dependencies of the system at homeostatic equilibrium. The replication of TTX-induced homeostatic responses has been treated more rigorously, with a more careful analysis of the relationship between the underlying model timescales and the experimentally observed scaling timescale. A new section has been added to establish a formal connection between the framework and the concept of metaplasticity. The analysis of dendritic compartmentalisation has been extended by incorporating local synaptic competition with size-dependent constraints, showing that this produces synaptic weight distributions that more closely resemble those observed experimentally. The replication of layer-specific local scaling has been extended with control simulations that identify the specific model components responsible for reproducing the experimental observations. These extensions have enabled us to derive six experimentally testable predictions that directly connect the theoretical framework to measurable quantities at the molecular timescale, synaptic weight distributions, and neuronal activity dynamics. Finally, the network model included in the previous version has been removed to focus more sharply on the single-neuron framework and its analytical results.

