## Supplementary Figures for "Local synaptic competition and global homeostatic regulation of synaptic resources as a unified framework for synaptic scaling"

### S1 Supplementary figures

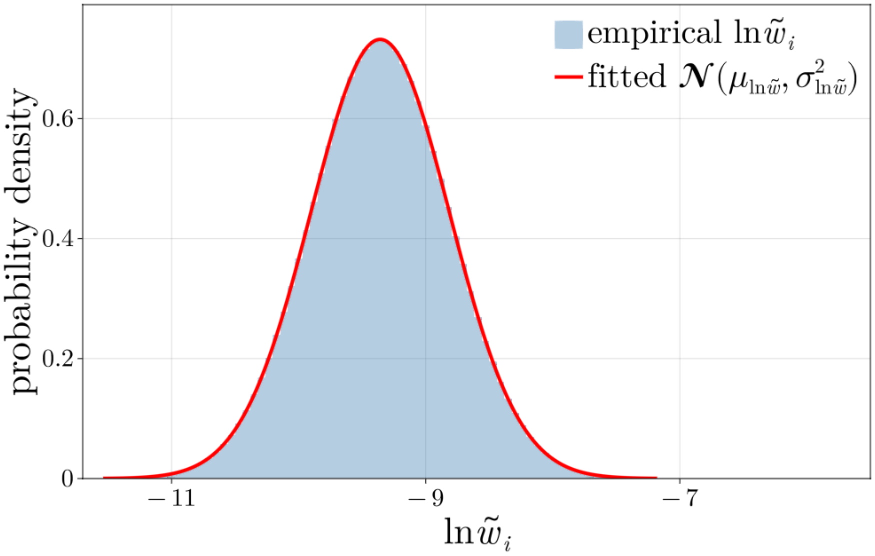


Figure 1: **Evaluation of lognormality.** The empirical distribution (blue) derived using a Monte Carlo process compared to a fitted lognormal distribution (red) with the same mean and variance as the empirical.

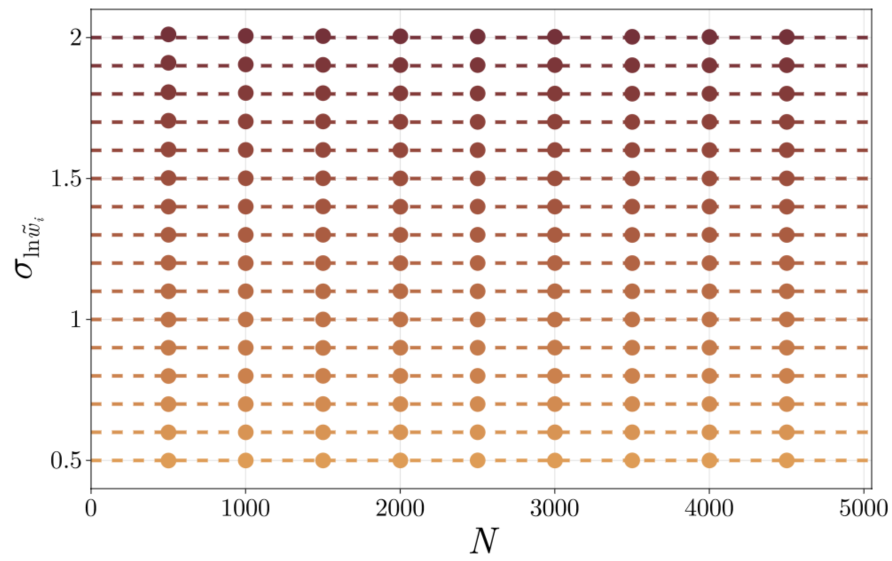


Figure 2: **Preservation of standard deviation for large $N$.** The log standard deviation of the empirical distribution of normalised weights, $\tilde{w}_i$, (dots) is similar to that of the effective weights, $w_i$, (dotted lines) for large values of the number of synapses, $N$.

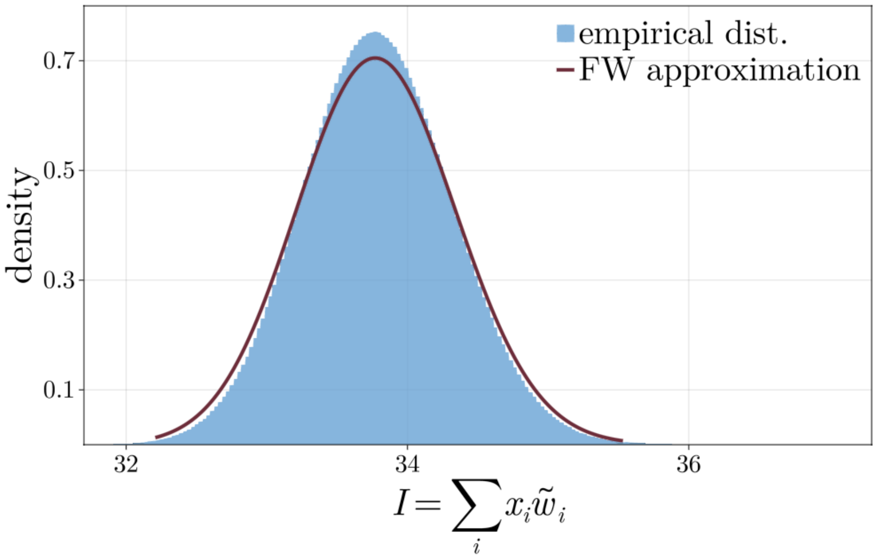


Figure 3: **Evaluation of input-drive approximation**. Fenton-Wilkinson approximation (red line) versus an empirical distribution (blue) derived using a Monte Carlo method. Estimates were obtained from 10.000 samples of $\tilde{w}_i$ and $x_i$ ($N = 10.000$), with 10.000 random assignments per sample.

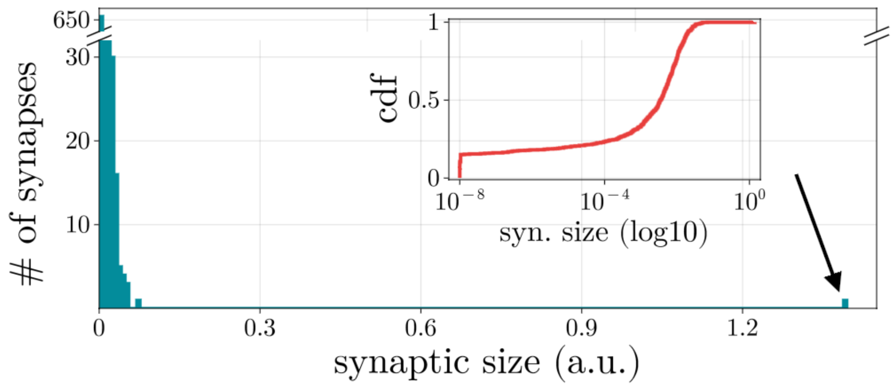


Figure 4: **Synaptic distribution under baseline conditions for additive STDP.** The example distribution here is dominated by a single overly potentiated synapse (black arrow) while the remaining synapses accumulate near the low hard bound, set to prevent negative or zero weights. Inset: cumulative distribution functions of the same distribution.
